# Bioelectric state transitions enable *de novo* feather bud formation in developing skin

**DOI:** 10.64898/2026.06.28.735123

**Authors:** Hans I-Chen Harn, Zhou Yu, Chih-Han Huang, Randle B. Widelitz, Ping Wu, Cheng-Ming Chuong, Robert Hsiu-Ping Chow

## Abstract

Tissue patterning is integral to development and regeneration, yet the factors that initiate morphogenetic patterning remain to be explored. Here, using embryonic chicken skin as a model, we show that perturbation of calcium signaling induces *de novo* feather bud formation in regions that normally do not form feather buds. This is achieved through coordinated changes in calcium dynamics, endogenous bioelectric currents, transcriptional regulation of calcium and potassium channel genes, and morphogen signaling. Different combinations of channel perturbations altered the number, distribution, size, and shape of induced feather buds. Live calcium imaging and extracellular electrophysiological recordings revealed homeostatic regulation, in which initially depressed calcium activity is followed by elevated calcium activity. Inward bioelectric currents emerge as *de novo* feather buds appear. Potassium channel blockade suppressed calcium dynamics, abolished endogenous currents, and inhibited new bud formation. Canonical feather morphogenesis pathways including Shh and β-catenin are induced in these new buds. Our findings support a model in which developmental bioelectricity contributes to regulating the threshold of feather bud formation. These results identify developmental bioelectricity as an unrecognized regulatory layer of tissue patterning that warrants further study.

**Bullet points:**

- Calcium signaling perturbation induces *de novo* feather bud formation in apteric skin
- Ion channel perturbations regulate the formation, distribution and shape of new buds across a continuum, depending on channel type(s) and perturbation strength.
- Elevated calcium activity and inward bioelectric currents accompany feather bud induction
- Developmental bioelectricity represents an unrecognized regulatory layer for morphogenesis

## 1. Introduction

Tissue patterning is a fundamental process during embryonic development. Tissue competence refers to the ability of cells or tissues to respond to morphogenetic cues and initiate morphogenesis, as exemplified in the formation of skin appendage buds. In most adult tissues, this developmental competence is lost and can only be regained under specialized regenerative conditions, such as axolotl limb regeneration (Tanaka, 2016) or wound-induced hair neogenesis (Harn et al., 2023; Ito et al., 2007). Understanding how to restore tissue competence and reinitiate morphogenesis is therefore a central question in regenerative biology. Feather bud formation in avian skin has served as a classic model for studying tissue competence, pattern formation and morphogenesis (Jiang et al., 2004; Olivera-Martinez et al., 2004). During skin development, feather-forming regions are organized into discrete patches, known as feather tracts (pterylae), which are separated by featherless or sparsely feathered regions called apteria (Lucas and Stettenheim, 1972). Within feather tracts, regularly spaced feather primordia emerge through coordinated epithelial–mesenchymal interactions, often described through reaction–diffusion mechanisms and morphogen gradients that generate periodic structures across the skin (Chuong et al., 2013; Painter et al., 2018). More recent work has expanded this framework to include dynamic processes such as propagating morphogenetic waves, which coordinate the temporal and spatial emergence of feather buds across the developing field (Ho et al., 2019; Inaba et al., 2019). However, the apteric region is not necessarily incapable of feather formation and experimental maneuvers can change the boundary between tract and apteric regions in developing embryos (Olivera-Martinez et al., 2004), thus, developing chicken skin provides an excellent model to study tissue competence.

Accumulating evidence suggests that competence may reflect a dynamic and emergent state of the tissue. For example, only subsets of cells respond to propagating inductive signals during morphogenetic wave progression, indicating that competence is temporally regulated (Ho et al., 2019; Inaba et al., 2019; Plikus et al., 2009). In addition, recent studies have demonstrated that physical properties such as cell density and tissue stiffness contribute to defining competent regions, implicating mechanical and collective cellular states as key regulators of morphogenesis (Harn et al., 2021; Ho et al., 2019; Jiang et al., 1999). These findings suggest that tissue competence arises from the integration of molecular, mechanical, and collective properties, yet the mechanisms remain to be investigated.

One emerging candidate for regulating tissue-level states is bioelectric signaling. Here, bioelectric state is used to describe the integrated electrophysiological properties of the tissue, including ion channel expression, channel activity, intracellular calcium dynamics, and endogenous ionic currents. Endogenous bioelectric cues, including ion fluxes and membrane potential differences, have been increasingly recognized as instructive regulators of development, regeneration, and large-scale pattern formation (Levin, 2021; Levin et al., 2017; McLaughlin and Levin, 2018). Bioelectric signals can act over long ranges, integrate cellular activity across tissues, and influence processes such as proliferation, migration, and differentiation (Jiang et al., 2021; Levin, 2021; McLaughlin and Levin, 2018). In diverse systems, modulation of ion channel activity has been shown to alter patterning outcomes, regenerate complex structures, and establish signaling thresholds that control cell fate decisions (Adams and Levin, 2013; Adams et al., 2007; Levin, 2007, 2013). Blocking gap junctions alters the periodic feather pattern formation (Tseng et al., 2024). Among these channel activities, calcium signaling represents a central integrator linking bioelectric activity to intracellular signaling networks. Calcium influx is tightly regulated by ion channels and can integrate signals through changes in amplitude and temporal dynamics, enabling precise control of downstream pathways. (Brodskiy and Zartman, 2018; Clapham, 2007; Webb and Miller, 2003). In developing tissues, calcium signaling has been implicated in coordinating cell migration, proliferation, and differentiation, and can interface with key morphogen pathways, including Sonic hedgehog (Shh) and Wnt/β-catenin signaling (Brodskiy et al., 2019; Li et al., 2020).

In the context of feather morphogenesis, calcium oscillations have been shown to coordinate mesenchymal cell behavior and contribute to feather elongation (Li et al., 2018). More recently, we showed that suppression of potassium channel activity converts spot-like feather primordia into stripe-like patterns during periodic feather patterning (Zitting et al., 2026). These observations raise the possibility that calcium-dependent bioelectric states may contribute to tissue competence. Here, we test whether perturbation of bioelectric signaling can induce *de novo* feather bud formation in apteric skin and how changes in ion channel activity, calcium dynamics, endogenous currents, and morphogen signaling accompany this process. Indeed, we found robust formation of new feather buds and studied the associated bioelectric changes. Our findings identify bioelectric state transitions as an unrecognized regulatory layer that gates morphogenetic competence and may be used to initiate collective tissue patterning.

## Results

### Calcium signaling perturbation induces *de novo* feather bud formation

To investigate whether calcium signaling regulates tissue competence during feather morphogenesis, we examined the effects of pharmacological perturbation of calcium-associated pathways in E9+3d chicken skin explants. Under normal conditions, apteric regions were devoid of feather buds, consistent with their normally non-competent state (Figure 1A, B). In contrast, inhibition of calmodulin signaling using KN62 induced robust *de novo* feather bud formation within the apteric skin at 2 and 5 µM (Figure 1B). Similar ectopic bud formation was observed following perturbation of T-type calcium channel blocker ethosuximide (ETH), L-type calcium channel blocker Nifedipine (NF) and the intracellular calcium chelator BAPTA-AM, indicating that disruption of calcium homeostasis can activate bud formation in normally non-bud-forming tissue (Figure 1B). Quantification of induced structures revealed a significant increase in apteric buds per unit area across calcium perturbation conditions compared to controls (Figure 1B). These new buds exhibited organized morphology resembling endogenous feather primordia, suggesting that calcium signaling perturbation does not merely trigger local cellular aggregation, but instead activates a coordinated morphogenetic program.

**Figure 1.**
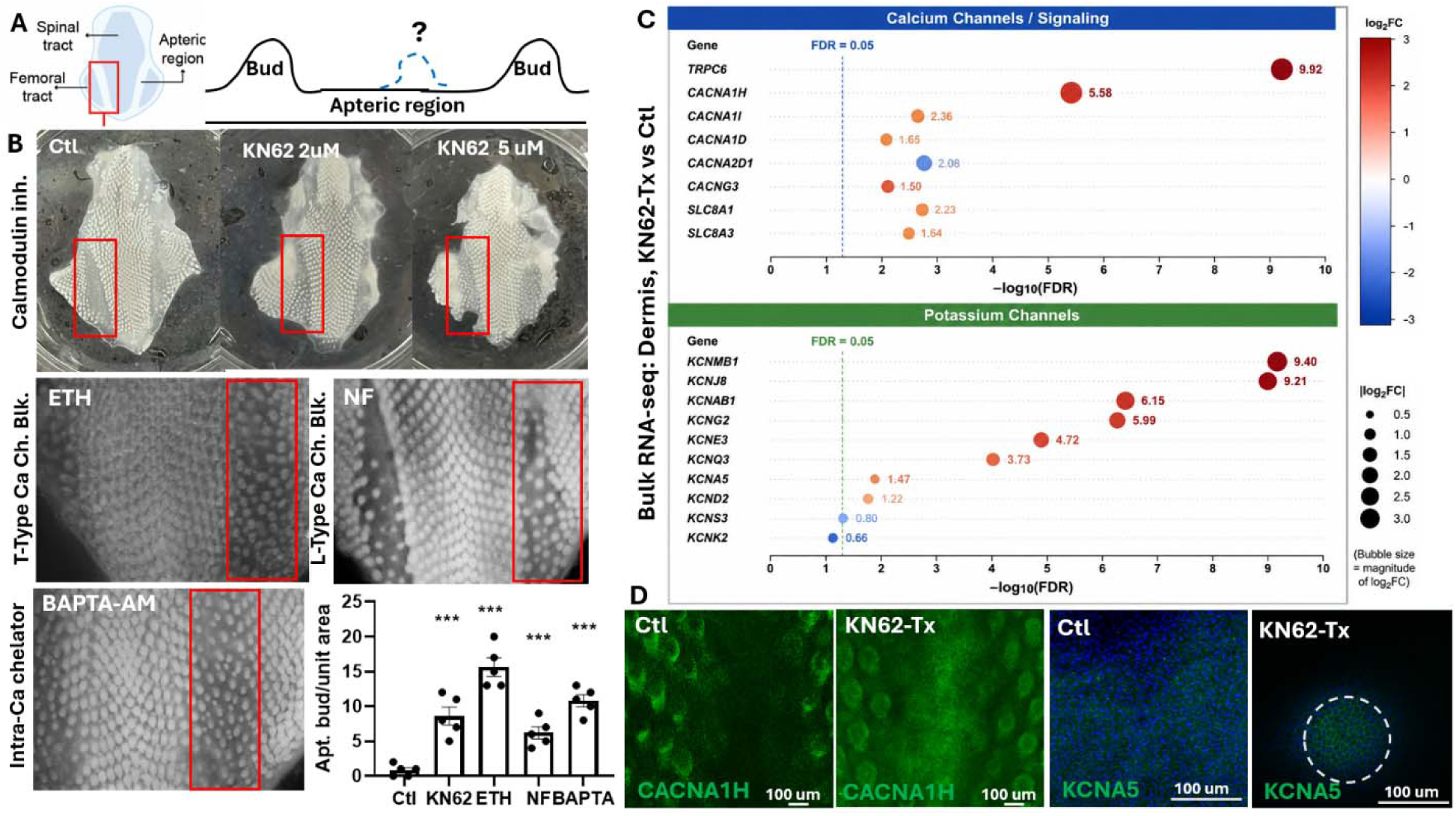
Calcium signaling perturbation induces de novo feather bud formation from apteric region. A. Schematic of embryonic chicken skin highlighting feather tract and apteric regions, with experimental focus on the apteric region that normally lacks feather buds. B. Brightfield images of E9+3d apteric skin explants treated with calmodulin inhibitor (KN62; 2 µM and 5 µM), showing induction of de novo feather buds compared to control. Perturbation of T-type calcium channel (ETH), L-type calcium channel (NF), and intracellular calcium chelation (BAPTA-AM), similarly induce ectopic bud formation. Red boxes indicate representative regions of bud induction. C. Bulk RNA-seq analysis of E9+3d dermal tissue comparing KN62-treated and control samples reveals differential expression of ion channel genes, including upregulation of calcium (e.g., CACNA1H) and potassium channel families. D. Induced ion channel gene expression. Immunofluorescence staining for T-type calcium channel CACNA1H and potassium channel KCNA5 in control and KN62-treated E9+3d samples, confirming increased expression in the apteric region following calcium signaling perturbation.

### Calcium signaling perturbation induces ion channel gene expression

To identify molecular changes associated with feather bud induction, we performed bulk RNA-seq on dermal tissue isolated from E9+3d KN62-treated and control explants. Differential expression analysis revealed broad upregulation of ion channel-associated genes following calcium signaling perturbation (Figure 1C). Notably, multiple calcium and potassium channel genes were upregulated in treated samples, including the T-type calcium channel gene CACNA1H, suggesting ion channel upregulation occurred in compensatory response to calcium pathway inhibition. Immunostaining further confirmed elevated CACNA1H and KCNA5 expression in the apteric region of KN62-treated tissue relative to controls (Figure 1D). Additionally, L-type calcium channel CACNA1G expression is also elevated in the apteric bud, although to a lesser extent (Figure 1C, SI Fig 1).

Together, these findings suggest that calcium pathway perturbation triggers compensatory ion channel upregulation, including upregulation of CACNA1H, and establish a potential mechanism that functions as a modulator of feather bud induction during morphogenesis.

### Calcium and potassium channel interactions quantitatively regulate the formation of new buds and the morphology of the resulting buds

To determine how different ion channels contribute to *de novo* feather bud formation, we next examined the effects of combinatorial perturbations targeting calcium and potassium signaling during *de novo* feather bud induction. Apteric skin explants were treated with inhibitors of calmodulin signaling (KN62), T-type calcium channels (ETH), intracellular calcium activity (BAPTA-AM), and potassium channels (4-AP), either alone or in combination (Figure 2A). While individual perturbations induced *de novo* feather buds, combinatorial perturbations generated distinct patterning outcomes characterized by differences in bud density, spatial distribution, size, and morphology. These perturbations produced a continuous range of feather bud density, size, and morphology, indicating that ion channel activity quantitatively modulates feather bud patterning. (Figure 2A).

**Figure 2.**
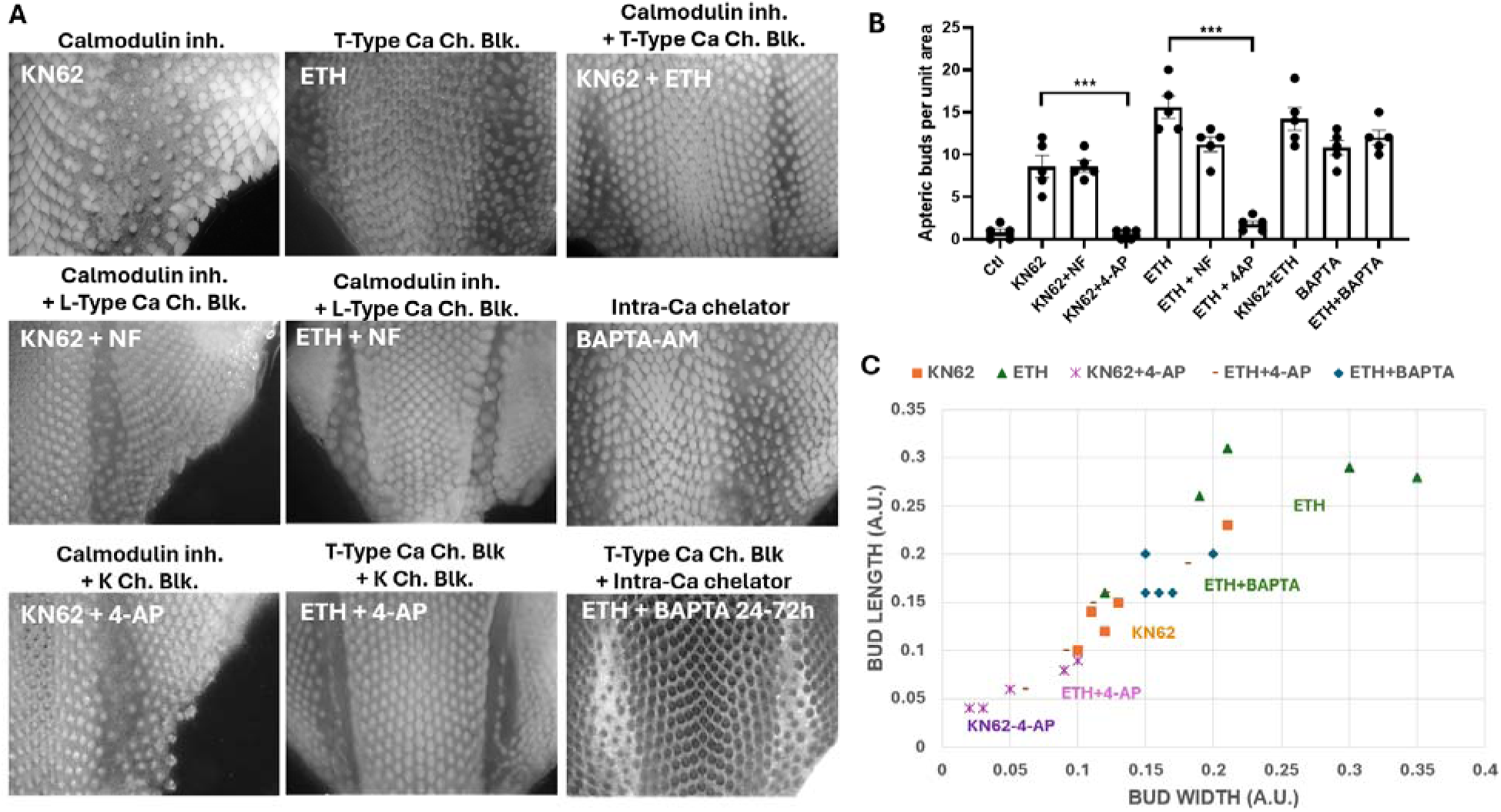
Calcium and potassium channel interactions to regulate the formation o new buds in apteric region and also the shape (aspect ration) of feather buds. A. Representative brightfield images of apteric skin explants treated with combinatorial perturbations of calcium signaling (KN62, ETH, NF and BAPTA-AM) and potassium channel inhibition (4-AP). Distinct patterns of feather bud formation are observed across conditions, demonstrating variable levels of tissue competence. B. Quantification of apteric bud density (buds per unit area) across treatment conditions. Combinatorial perturbations produce graded effects, with certain combinations enhancing or suppressing bud formation relative to single treatments. C. Scatter plot of bud length versus width across conditions, illustrating continuous variation in bud morphology. Data points cluster according to treatment, revealing that ion channel interactions regulate both the formation and morphology of new buds.

Morphological analysis revealed substantial variation in feather bud density, size, and morphology according to treatment conditions (Figure 2A-C). Certain treatments produced relatively large and elongated buds, whereas others generated smaller or less organized structures, indicating that ion channel activity quantitatively and qualitatively influences both bud initiation and progression of bud morphogenesis. Either KN62 or ETH treatment alone induced robust ectopic buds within apteric regions. Potassium channel inhibitor 4-AP combined with either KN62 or ETH reduced the number and size of ectopic buds. Adding BAPTA-AM to ETH reduced length and width of *de novo* buds (Figure 2C). Interestingly, combination of T- and L-type calcium channel inhibitor (ETH and NF) led to enlarged buds (Figure 2A, C), implying T- and L-type calcium channels play different roles in the size control of feather bud development. Quantification of apteric buds per unit area demonstrated that different combinations of calcium and potassium pathway perturbations generated distinct levels of inductive response (Figure 2B). Scatter plot analysis of bud length versus width further revealed that induced structures occupied a continuous morphological spectrum rather than discrete states (Figure 2C).

Notably, potassium channel inhibition exerted a strong modulatory effect on calcium-induced morphogenesis. Co-treatment with 4-AP markedly attenuated feather bud induction across multiple calcium perturbation conditions, supporting a functional interaction between potassium conductance and calcium-dependent pattern formation. These findings suggest that feather bud formation depends on a coordinated bioelectric state established through coupled calcium and potassium channel activity. Blocking potassium channels can cause the cell membrane potential to remain depolarised, which in turn inactivates T-type calcium channels (Cain and Snutch, 2010) and reduces the driving force for calcium entry through L-type calcium channels. Correspondingly, CACNA1H expression is also upregulated in the apteric buds induced by calcium signaling blockage (ETH, BAPTA, ETH+NF), and downregulated where *de novo* buds are abrogated (ETH+4-AP or ETH+BAPTA) (SI Fig 2), suggesting an important role in mediating calcium signaling in this process.

Together, these results suggest that coordinated calcium and potassium signaling contributes to establishing a permissive bioelectric state for feather bud formation, while distinct ion channel perturbations quantitatively regulate the subsequent distribution, size, and morphology of the developing feather buds.

### Homeostatic regulation of ion channel activities reshapes calcium dynamics and bioelectric currents

Because perturbation of calcium and potassium signaling quantitatively altered feather bud induction, we next asked whether these treatments modify the underlying bioelectric current of the tissue. To monitor calcium dynamics during feather bud formation, E9 skin explants expressing the calcium reporter RCAS-mCherry-GCaMP6 were imaged over time, and the feather tract, interbud, and apteric regions were analysed (Figure 3A). Under control conditions, endogenous feather buds exhibited elevated and spatially organized calcium activity compared to adjacent interbud regions, indicating that calcium activity is associated with the growth of feather buds (Figure 3B).

**Figure 3.**
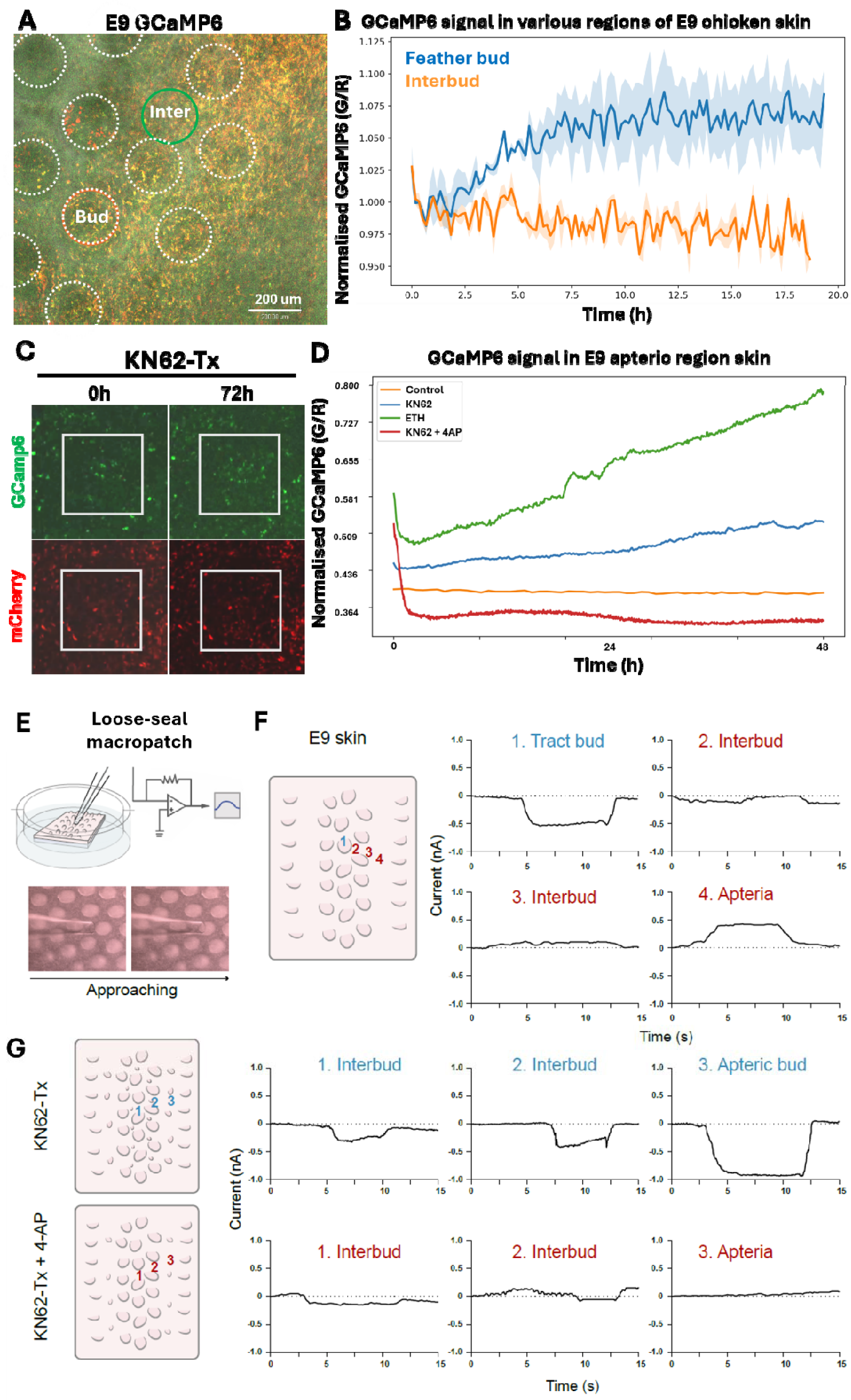
Ion channel–driven calcium dynamics reshape bioelectric currents to establish tissue competence. A. Live imaging of mCherry-GCaMP6 activity in E9 chicken skin explant, showing spatial differences between feather bud and interbud regions. Dotted circle: feather bud. Green circle: representative region of interest for interbud GCaMP6 signal quantification. B. Quantification of normalized GCaMP6 fluorescence over time reveals distinct calcium dynamics in bud versus interbud regions, with rising and maintaining elevation associated with bud formation. Shadow: SEM. C. Time-course imaging of GCaMP6 and mCherry in control and KN62-treated explants over 72 hours, demonstrating altered calcium signaling dynamics following perturbation. D. Quantitative analysis of calcium activity over time across multiple treatment conditions (control, KN62, ETH, KN62 + 4-AP), showing treatment-specific trajectories of calcium signaling. E. Schematic of electrophysiological recording setup for measuring bioelectric currents in skin explants using a loose-seal macropatch. F. Representative recordings of bioelectric currents across different tissue regions (tract bud, interbud, apteric region) in E9 skin. E. Representative recordings of bioelectric currents in KN62 and KN62+4-AP treatment conditions. Perturbations alter both the magnitude and polarity of currents, with transitions toward inward currents associated with induced competence.

Following calcium pathway perturbation, Apteric regions undergoing *de novo* feather bud induction showed an initial decrease in calcium activity, before progressively recovering and exceeding baseline levels (Figure 3C, D). Notably, this delayed increase is consistent with the compensatory transcription of ion channels observed in Figure 1 and suggests that elevated calcium activity arises as an adaptive response (Figure 3C). Temporal analysis revealed that calcium activity decreases initially upon calcium perturbation (KN62 and ETH) in the first hour and then progressively increased over the remaining 47 hours. In contrast, KN62+4-AP treatment, where no *de novo* bud would form, yielded a much steeper drop in GCaMP6 signal, and remained low throughout the recording. This suggests that blocking potassium channels with 4-AP also prevented the membrane potential from returning toward its normal negative resting level (Faber and Sah, 2003). These observations support a model in which an initial reduction in calcium signaling triggers homeostatic and transcription regulation of ion channels, leading to eventual elevated calcium activity that enables new bud formation (Figure 3D).

To determine whether altered calcium activity was associated with changes in tissue-level bioelectric signaling, we next measured endogenous ionic currents using a loose-seal macropatch (Hilgemann and Lu, 1998) to record extracellular bioelectric currents across different skin regions and treatment conditions (Figure 3E). E9 feather tract buds displayed characteristic inward currents distinct from neighboring interbud, and apteric regions showed outward currents (Figure 3F). Importantly, calcium signaling perturbation induced substantial changes in current profiles within the interbud and especially the apteric region, shifting the normally outward currents of apteric tissue toward inward current states characteristic of endogenous feather buds. (Figure 3G). The emergence of inward currents preceded *de novo* bud formation and therefore represents an early bioelectric current signature of feather bud induction. Potassium channel inhibition with 4-AP strongly disrupted this transition. Co-treatment with 4-AP markedly reduced calcium activity and nearly abolished detectable bioelectric currents in KN62-treated explants, coinciding with suppression of *de novo* feather bud formation (Figure 3G). These findings indicate that coordinated potassium conductance is required to sustain the calcium-dependent bioelectric state associated with bud formation and its graded morphology.

Together, these results demonstrate that ion channel perturbation remodels both calcium dynamics and endogenous bioelectric currents during feather bud induction. The emergence of elevated calcium activity and inward current states in apteric tissue suggests that induction of morphogenesis is associated with a distinct bioelectric state that facilitates *de novo* feather bud formation.

### Bioelectric tuning of morphogen signaling

To determine whether bioelectric state transitions engage canonical morphogen pathways during the formation of new feather buds, we examined the expression of signaling components associated with feather bud initiation following calcium signaling perturbation. RNA-seq analysis of E9+3d KN62-treated apteric epidermis showed upregulation of Shh and canonical Wnt programs compared to control (Figure 4A). Wholemount immunostaining also showed that KN62-treated apteric skin undergoing *de novo* bud formation displayed localized activation of canonical feather morphogen pathways, including Sonic hedgehog (Shh) and β-catenin signaling (Figure 4B). Apteric feather buds also showed a translation of calmodulin in the nucleus (SI Fig 3). Additionally, Shh and β-catenin signals were also elevated in the KN62-treated feather tract buds, resulting in small bud-like protrusions (Figure 4B). These findings indicate that perturbation of calcium-dependent bioelectric states is associated with activation of canonical morphogen pathways in tissue that is normally non-competent for appendage formation.

**Figure 4.**
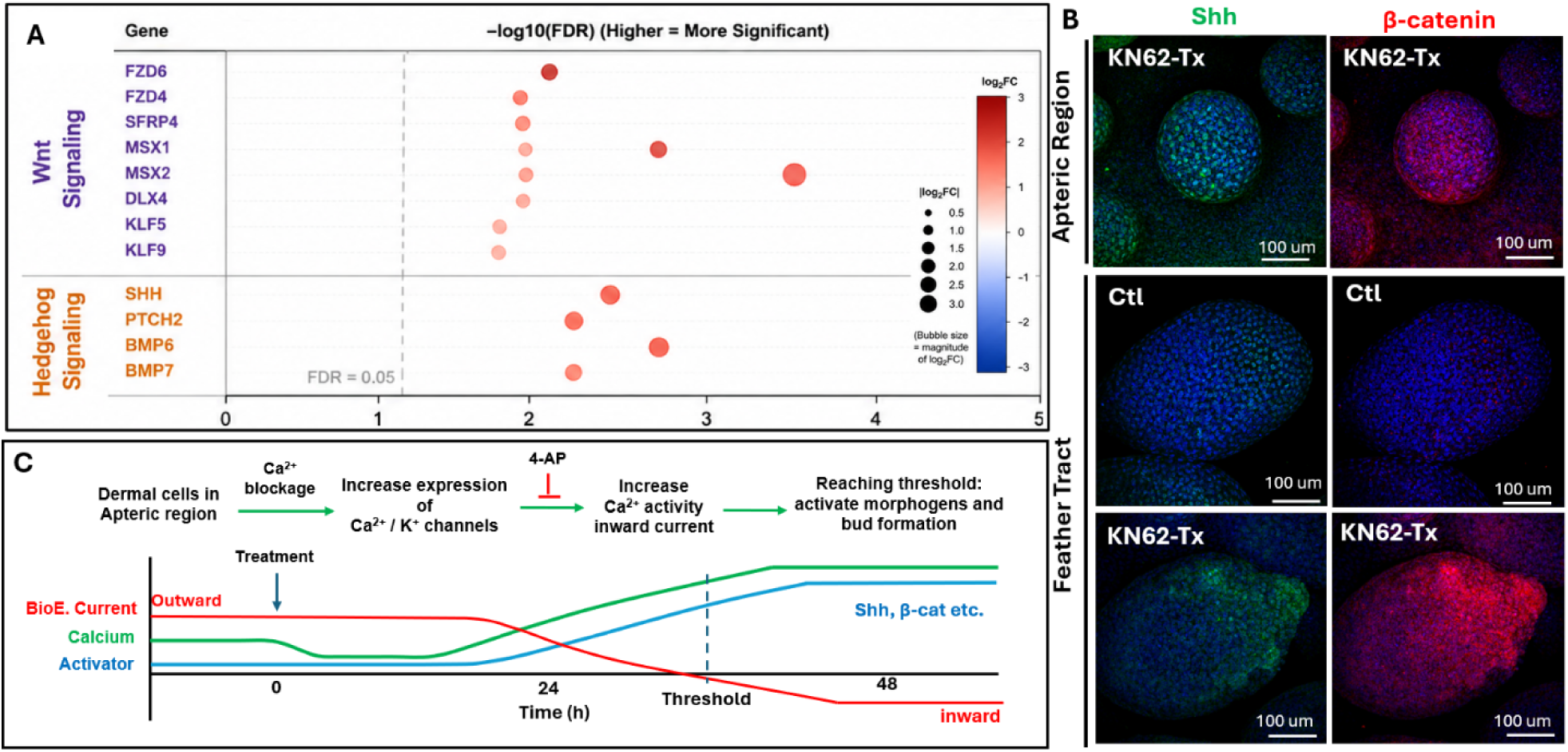
Bioelectric reprogramming activates morphogen signaling to initiate de novo feather bud formation. A. RNA-seq analysis of apteric epidermis showing Wnt and Hedgehog signaling associated genes are upregulated in KN62-Tx samples in comparison to that of control. B. Immunostaining for morphogen pathway components Sonic hedgehog (Shh) and β-catenin in apteric regions and feather tracts under control and KN62-treated conditions. These canonical morphogens are activated in the apteric buds and even in the mature tract buds, where smaller additional buds are observed on native tract buds. C. Proposed model of bioelectric reprogramming observed in this study: calcium signaling perturbation induces compensatory upregulation of calcium and potassium channels, leading to increased calcium activity and a shift toward inward bioelectric currents. Once a threshold is reached, morphogen signaling pathways (e.g., Shh, β-catenin) are activated, initiating de novo feather bud formation.

Because calcium and potassium channel perturbations altered both calcium dynamics and endogenous ionic currents (Figure 3), we next integrated these findings into a mechanistic model linking ion channel remodeling to morphogen activation (Figure 4C). Under normal condition, the apteric skin maintains a moderate level of intracellular calcium concentration, outward bioelectric current and low activator (morphogen) activity. Pharmacological inhibition of calcium signaling leads to a drop in calcium activity, which then induces compensatory upregulation of calcium and potassium channels, a process that takes around 24 hours. The elevated ion channel expression in turn increase calcium dynamics and a transition toward inward bioelectric currents within apteric tissue. Once inward bioelectric currents and calcium activity reached a permissive bioelectric state, canonical morphogen pathways including Shh and β-catenin became activated, allowing feather bud morphogenesis to proceed 48 hours after the initial treatment. Notably, conditions that disrupted the bioelectric transition, including potassium channel blockade with 4-AP, strongly suppressed feather bud induction and reduced calcium activity and tissue currents, supporting the requirement for a coordinated bioelectric state that facilitates morphogen activation. Together, these findings suggest that bioelectric state transition functions as a previously unrecognized regulatory layer that gates morphogenetic competence and enables *de novo* feather bud formation in the originally apteric region of embryonic skin.

## 3. Discussion

In this study, we demonstrate that perturbation of calcium-dependent bioelectric signaling induces *de novo* feather bud formation in normally apteric skin. This developmental response is accompanied by coordinated changes in ion-channel expression, calcium dynamics, and endogenous bioelectric currents, and canonical morphogen signaling. How is this achieved? One possible explanation is that apteric tissue normally fails to achieve the activation threshold required to initiate the new feather bud formation. A concept emerging from this work is that morphogenesis may require tissues to surpass a critical activation threshold so appendage formation can be initiated and that altering bioelectric state of the field can tune the threshold. Turing-type models propose that periodic pattern formation emerges through the balance between local activation and long-range inhibition, in which activating signals must overcome inhibitory constraints to trigger morphogenesis (Perrimon et al., 2012).

Perturbation of calcium signaling triggered homeostatic regulation of ion channel activities and transcriptional regulation of channel genes, thus reprogramming the bioelectric landscape and leading to inward bioelectric currents. Our findings identify compensatory transcriptional regulation of ion channel genes as a potential mechanism underlying the transition toward a competence-permissive state. Pharmacological suppression of calcium signaling unexpectedly induced transcriptional upregulation of calcium and potassium channel families, including increased expression of the T-type calcium channel CACNA1H (Fig. 1C, D). Whether this transcriptional response promotes morphogenesis through altered channel activity, downstream transcriptional programs, or modulation of morphogen production remains an important question for future investigation. Rather than simply reducing calcium activity, calcium pathway inhibition instead triggered an adaptive bioelectric response that was associated with elevated calcium dynamics and inward bioelectric currents within apteric tissue (Fig. 3). Notably, these current transitions were localized to feather-forming regions and emerged during detectable morphogenesis, supporting the idea that bioelectric signaling acts locally within competent tissue domains. We propose that this compensatory remodeling progressively raises endogenous bioelectric activity toward a state that favors periodic feather bud formation. Once this state is reached, downstream developmental programs including Shh and β-catenin signaling are activated, enabling *de novo* feather bud formation to proceed (Fig. 4B). In this framework, bioelectric currents may contribute to feather bud formation through interactions with canonical morphogen pathways.

These results suggest that the impact of ion-channel activity on feather morphogenesis is regulated at two distinct levels. *First,* the initiation of *de novo* feather bud formation appears to require a permissive bioelectric state, consistent with an activation threshold for morphogenesis. Conditions that failed to establish elevated calcium activity and inward bioelectric currents, such as potassium channel blockade with 4-AP, did not produce feather buds, whereas perturbations that generated these bioelectric changes induced bud formation. *Second*, once feather bud formation was initiated, different ion channel perturbations produced variation in the number, spacing, size and shape of buds (Fig. 2B, C), indicating that the morphogenetic response itself is quantitatively tunable rather than binary. This two-level model is consistent with previous studies of skin patterning. Mesenchymal cell density has been proposed to reach a critical level before feather tract formation can begin (Ho et al., 2019), while cell shape anisotropy influences the fidelity of periodic feather patterning once buds are established (Curantz et al., 2022). Mechanistically, calcium signaling is well positioned to coordinate such integration because it links membrane bioelectric currents with intracellular signaling pathways and transcriptional regulation (Levin, 2021). Calcium dynamics can regulate cellular responses through changes in signal amplitude and temporal oscillations, enabling cells to integrate multiple environmental and developmental cues (McLaughlin and Levin, 2018). Previous studies demonstrated that calcium oscillations coordinate mesenchymal cell behavior during feather elongation (Li et al., 2018), while the findings here suggest that elevated calcium activity also contributes to the earlier transition into a competence-permissive state (Fig 4B). Inward bioelectric currents therefore emerge as a candidate that contributes to tuning morphogenetic thresholds.

Several limitations should be considered. Future studies should determine whether basal calcium activity differs between feather-forming and apteric regions and how such differences contribute to morphogenetic competence. The present study relies primarily on pharmacological perturbations, and future genetic approaches will be important for defining the specific ion channel networks that regulate competence transitions. In addition, although our data support a compensatory ion channel remodeling response following calcium signaling suppression, the mechanisms linking reduced calcium signaling to transcriptional upregulation of calcium and potassium channel programs remain unclear. How coordinated ion channel activity elevates calcium dynamics and endogenous bioelectric currents to activate downstream morphogen pathways remains to be determined. Lastly, whether bioelectric signaling acts by altering tissue competence, inductive signaling, or the crosstalk between these processes remains an important question for future investigation.

In summary, we report the unexpected finding that modulating the bioelectric state of developing skin is sufficient to restore tissue competence in the apteric region, leading to the *de novo* formation of feather buds. Together, our findings suggest that tissue bioelectric states represent an additional regulatory layer contributing to tissue patterning and morphogenesis. This system could allow developing tissues to integrate molecular, mechanical, and collective cellular information into a unified decision-making process governing tissue patterning. Modulating membrane potential may provide a means to restore developmental competence in adult tissues, thereby enabling regenerative morphogenesis.

## Author contributions

**Hans I-Chen Harn**: Data curation, Formal analysis, Investigation, Methodology, Writing – original draft. **Zhou Yu**: Data curation, Formal analysis, Investigation, Methodology, Validation, Visualization, Writing – original draft. **Chih-Han Huang**: Data curation, Formal analysis, Investigation, Methodology, Validation. **Randall Widelitz**: Conceptualization, Funding acquisition, Supervision. **Ping Wu**: Conceptualization, Investigation, Validation. **Cheng-Ming Chuong**: Conceptualization, Funding acquisition, Resources, Supervision, Validation, Writing – original draft, Writing – review & editing. **Robert Hsiu-Ping Chow**: Conceptualization, Formal analysis, Funding acquisition, Methodology, Resources, Supervision, Validation, Writing – original draft, Writing – review & editing.

## Declaration of generative AI and AI-assisted technologies

ChatGPT Edu was used to polish grammatical errors. No generative AI tool was used.

## Data availability

## STAR Methods

### Resource Availability

#### Lead contact

Requests for further information and resources should be directed to and will be fulfilled by the lead contact, Hans Harn.

#### Materials availability

All unique/stable reagents generated in this study are available from the lead contact with a completed materials transfer agreement.

#### Data and code availability

RNA-seq data generated in this study have been deposited in [GEO/SRA accession number] and are publicly available as of the date of publication.

All custom analysis scripts used for image processing, calcium imaging quantification, RNA-seq analyses, and statistical analyses are available from the lead contact upon reasonable request.

Any additional information required to reanalyze the data reported in this paper is available from the lead contact upon request.

## Experimental models

### Avian embryos

Chicken embryos were harvested from pathogen-free fertilized White Leghorn eggs (SPAFAS, Preston, CT). Membrane-GFP transgenic quail eggs were kindly provided by the Lansford laboratory (Huss et al., 2015).

Embryos of both sexes were included randomly. Sex was not determined because embryos were analyzed at developmental stages where sex identification is technically difficult and unlikely to influence experimental outcomes.

### Chicken skin explant culture

Dorsal skin explants were dissected from chicken embryos at the indicated developmental stages in Hank’s Buffered Saline Solution (HBSS; Thermo Fisher, 14170161).

Explants were transferred onto cell culture inserts containing 0.4 μm pores (Falcon, 62406-163) and maintained at an air-liquid interface. Culture medium consisted of: DMEM (Thermo Fisher, 10569044), 10% fetal bovine serum (Thermo Fisher, A5670801), 2% chicken serum, 1% Penicillin-Streptomycin (Thermo Fisher, 15070063). Approximately 1 mL of medium was added beneath each insert. Explants were cultured at 37°C under 5% CO2 and 90% humidity.

## Methods

### Pharmacological treatments

Small-molecule inhibitors were dissolved in DMSO and diluted into culture medium to the indicated final concentrations.

Treatment conditions included:

- KN62 (calmodulin inhibition), (MedChemExpress, HY-13290)
- KN93 (calmodulin inhibition), (MedChemExpress, HY-15465)
- Ethosuximide (ETH, T-type calcium channel blocker), (Selleckchem, S4626)
- Nifedipine, (NF, L-type calcium channel blocker), (Selleckchem, S1808)
- BAPTA-AM (intracellular calcium chelator), (Selleckchem, S7534)
- 4-Aminopyridine, (4-AP, potassium channel blocker), (Sigma, 275875)
- Tetraethylammonium, (TEA, potassium channel blocker), (Sigma, T2265)
- ABT-639 (T-type calcium channel blocker), (Cayman Chemical, No. 28742)

Vehicle controls received equivalent DMSO concentrations.

### Histology

Explants were fixed overnight in 4% paraformaldehyde (PFA) at 4°C.

Samples were dehydrated through a graded ethanol series, cleared in xylene, embedded in paraffin, and sectioned at 6 μm thickness.

Hematoxylin and eosin staining was performed according to standard protocols.

For whole-mount analyses, tissues were fixed in 4% PFA and stored in PBS containing sodium azide at 4°C.

### Immunofluorescence staining

#### Antibodies

| Antibody | Host | Company | Catalog number | Dilution |
| --- | --- | --- | --- | --- |
| CACNA1H | Rabbit | Proteintech | 28092-1-AP | 1:50 (IHC), 1:200 (WM) |
| Calmodulin | Mouse | Biotech | NB120-5494-50ul | 1:50 (IHC), 1:200 (WM) |
| β-catenin | Mouse | Proteintech | 66379-1-Ig | 1:50 (IHC), 1:200 (WM) |
| Shh | Rabbit | Proteintech | 20697-1-AP | 1:50 (IHC), 1:200 (WM) |
| KCNA5 | Rabbit | BiCell | 20225 | 1:50 (IHC), 1:200 (WM) |
| Twist1 | Rabbit | Proteintech | 25465-1-AP | 1:50 (IHC), 1:200 (WM) |

#### Paraffin section immunofluorescence

Sections (IHC) were subjected to standard antigen retrieval and blocking procedures. Primary antibodies were incubated overnight at 4°C. Secondary antibodies were incubated for 1 hour at room temperature. Phalloidin and DAPI were included at 1:1000 dilution during secondary incubation.

#### Whole-mount immunostaining

Whole-mount (WM) tissues were permeabilized in methanol and incubated in 3% H2O2 for 30 minutes. Samples were blocked for 1 hour and incubated with primary antibodies overnight at 4°C with agitation. Following TBST washes, samples were incubated with secondary antibodies and Hoechst overnight at 4°C. Samples were cleared through graded glycerol solutions to 100% glycerol before imaging. Primary antibodies were typically used at 1:400 dilution.

### RNA isolation and library preparation

Skin explants were harvested and regions of interest were microdissected. Samples were incubated in calcium-magnesium-free solution containing 0.25% EDTA for 15–25 minutes on ice. Epidermis and dermis were mechanically separated under a dissecting microscope. Total RNA was isolated using the RNeasy Mini Kit (QIAGEN). RNA library preparation and sequencing were performed by Novogene Co., Ltd. (Sacramento, CA, USA). Sequencing libraries were sequenced on an Illumina NovaSeq 6000 platform using a paired-end 150-bp (PE150) configuration. An average of approximately 76 million raw reads was generated per sample, with >91% of bases achieving a quality score of Q30 or higher.

### RNA-seq analysis

Raw sequencing reads were quality controlled and filtered by Novogene Co., Ltd. (Sacramento, CA, USA) before downstream analyses. Reads were aligned to the Gallus gallus reference genome assembly galGal6 (UCSC) using the STAR aligner implemented in Partek Flow. Gene-level read counts were generated based on the corresponding genome annotation and normalized for downstream analyses. Principal component analysis (PCA) was performed on normalized gene expression data to assess sample clustering and global transcriptional variation. Differential expression analysis between KN62-treated and control samples was performed using the DESeq2 workflow implemented in Partek Flow. Genes with a false discovery rate (FDR) < 0.05 and an absolute log2 fold change > 1 were considered significantly differentially expressed. All analyses were performed using biologically independent replicates where n=2.

### RCAS-GCaMP6-mCherry virus production

The RCAS-GCaMP6s-T2A-mCherry construct was generated previously (Li et al., 2018). Replication-competent RCAS virus was produced and concentrated by ultracentrifugation (26,000 rpm, 1.5 h; Beckman Coulter L8-80M, SW28 rotor). Concentrated virus was used for infection of embryos and skin explants.

### Live calcium imaging

Live xyzt, temperature and humidity controlled imaging was performed according to Harn et al (Harn et al, EMBO J, 2026). Confocal microscope (Stellaris 5, Leica) with a temperature (37 oC) and humidity (5% CO2, 90% relative humidity) controlled chamber (Oko-Lab) and 10x air objective lens. Z-stacks (∼50 um) were collected at 10-min intervals for 48h. Laser power, gain, and exposure time were kept minimal and constant for all recording sessions.

Time-lapse image stacks were acquired simultaneously in the GCaMP6 (green) and mCherry (red) channels. The resultant tif files were saved and analyzed by selecting the respective region of interests, whether it’s interbud, bud, or de novo buds.

The green/red ratio was calculated as:

G/R = Mean GCaMP6 / Mean mCherry

For normalized analyses:

R(t)/R0

or

ΔR/R0

were calculated using baseline fluorescence values obtained from the initial recording period.

Raw G/R values and normalized traces displayed identical temporal trends.

For replicate analyses, traces were aligned onto a common normalized time axis and averaged.

For presentation purposes only, selected control traces were smoothed using a Savitzky–Golay filter. All quantitative analyses were performed on unsmoothed data.

Biological replicates represent independently cultured skin explants obtained from separate embryos.

### Loose-seal macropatch recordings

#### Pipette fabrication

Loose-seal macroelectrodes were fabricated as described previously (Hilgemann, 1998). Pipettes were pulled from borosilicate glass capillaries and fire-polished to approximately 0.1 mm inner tip diameter.

#### Recording setup

Recordings were performed using an Olympus MVX10 microscope equipped with a 1× MV PLAPO objective. An Ag/AgCl pellet served as the reference electrode. Signals were amplified using a Dagan PC-ONE patch-clamp amplifier. Currents were filtered at 100 Hz and digitized at 6.25 Hz. The loose-seal macropatch configuration measures extracellular ionic currents generated by local tissue activity and does not directly measure membrane potential. Current amplitudes were quantified over 1 second. Multiple tests were performed to ensure consistency.

## Quantification and statistical analysis

### Feather bud quantification

Whole-mount images were acquired using identical imaging parameters.

Measurements included: bud density (buds/mm²), bud length and bud width. Bud density was normalized to tissue area. Length and width were measured in Fiji/ImageJ using standardized anatomical landmarks. The length is defined as the distance between anterior and posterior end of a feather bud, while the width is the distance between the two lateral sides. Multiple fields were averaged to generate a single biological replicate. At least 3 biological replicates were used.

### Statistical analyses

Statistical analyses were performed using GraphPad Prism version 8.

Data are presented as mean ± SD unless otherwise indicated.

Statistical tests included:

- unpaired two-tailed Student’s t test
- one-way ANOVA
- two-way ANOVA
- Tukey’s post hoc test

Significance thresholds:

- p < 0.05 ** p < 0.01 *** p < 0.005 **** p < 0.001

Sample sizes (n) are indicated in figure legends.

Sample results outside of SD*2 were defined as outliers and excluded. Experiments were randomized. Investigators were blinded during quantitative image analyses but not during experimental procedures.

