## Supplementary Figures for "Bioelectric state transitions enable *de novo* feather bud formation in developing skin"

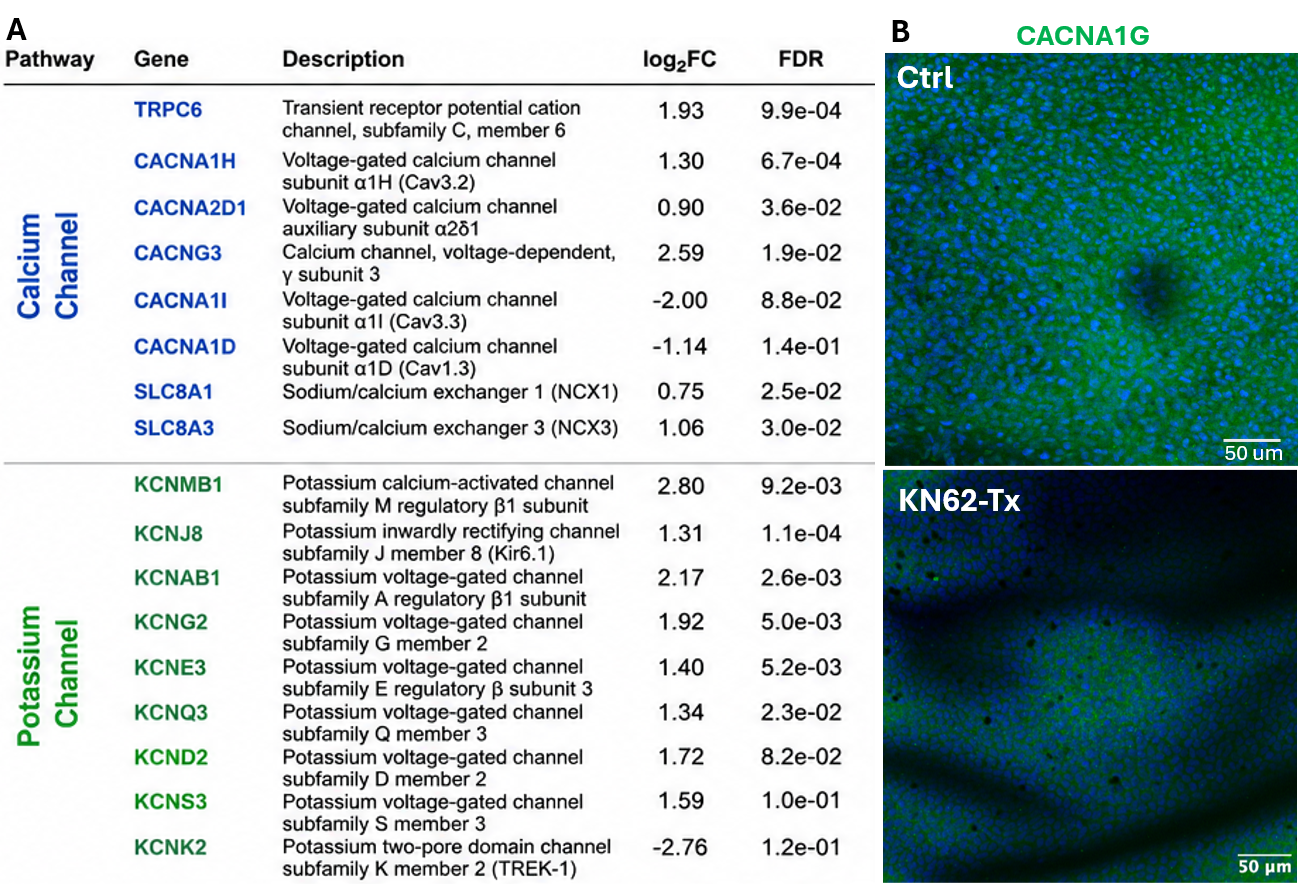
**SI Fig 1. T-type calcium channel CACNA1G is moderately expressed in the KN62-treatment induced apteric buds.**

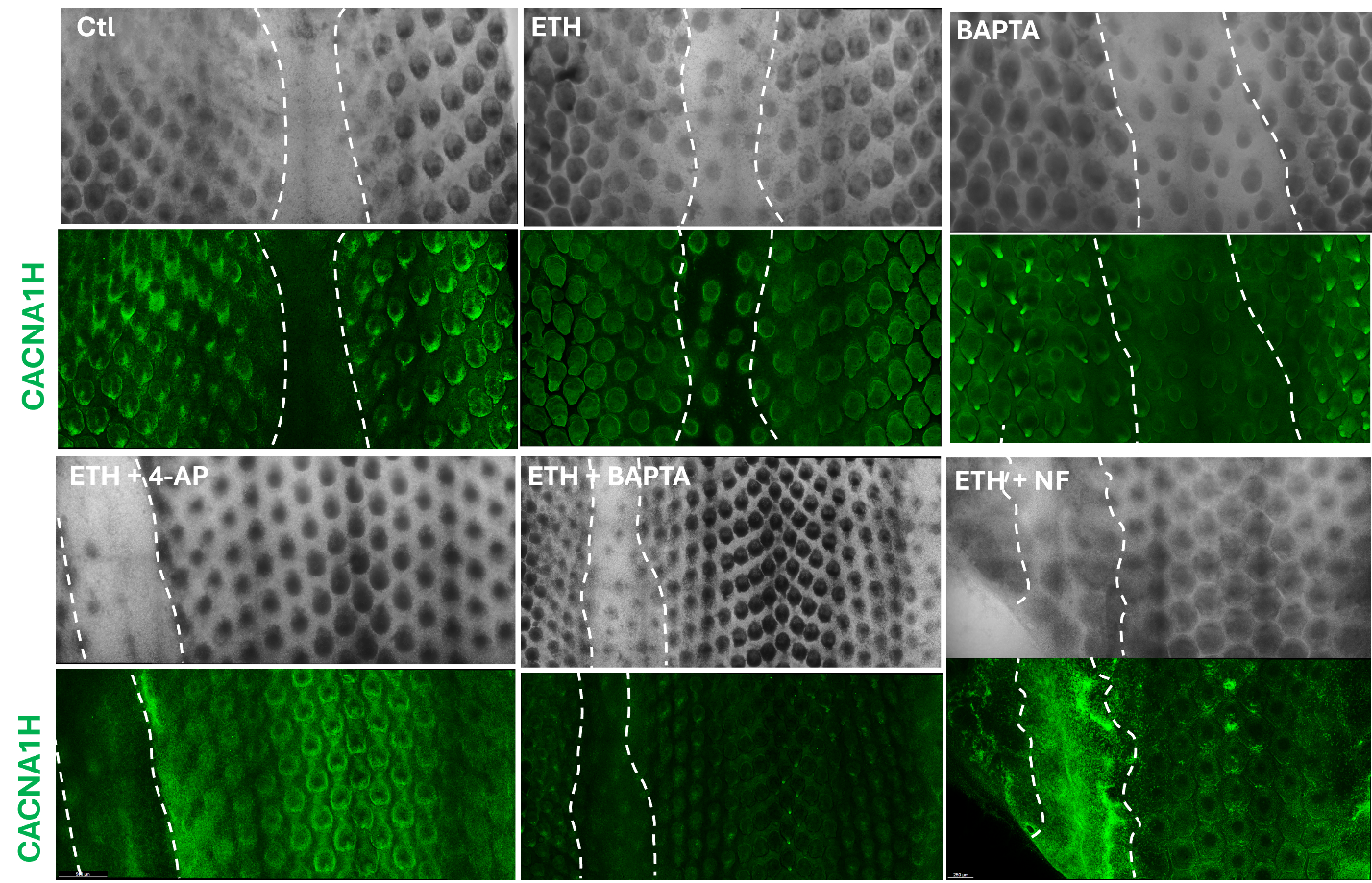

**SI Fig 2. CACNA1H is upregulated in the apteric buds induced by calcium signaling perturbation.**

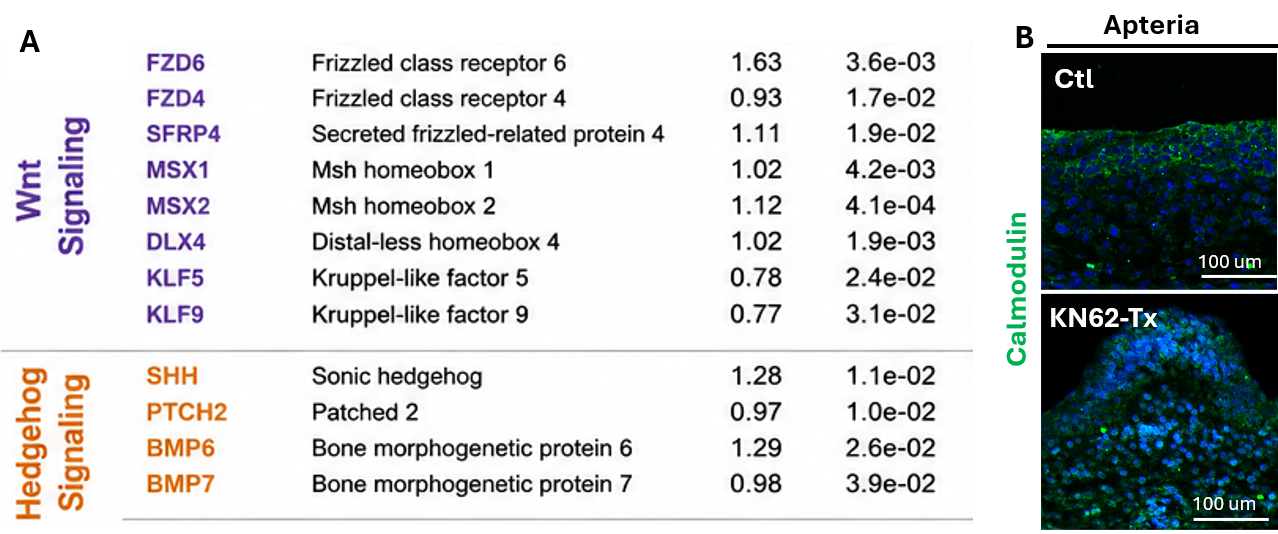

**SI Fig 3. KN62 treatment upregulated Wnt and Shh signaling in the apteric dermis and translocated calmodulin into nucleus in the apteria.**

A. Bulk-RNA-seq analysis of KN62-Tx vs control apteric dermis showing upregulation of genes associated with Wnt and Hedgehog signaling
B. KN62-Tx translocated calmodulin expression into the nucleus in the de novo apteric bud.
